# Population genetics of *Glossina palpalis gambiensis* in the sleeping sickness focus of Boffa (Guinea) before and after eight years of vector control: no effect of control despite a significant decrease of human exposure to the disease

**DOI:** 10.1101/2023.07.25.550445

**Authors:** Moise S. Kagbadouno, Modou Séré, Adeline Ségard, Abdoulaye Dansy Camara, Mamadou Camara, Bruno Bucheton, Jean-Mathieu Bart, Fabrice Courtin, Thierry de Meeûs, Sophie Ravel

## Abstract

Human African trypanosomosis (HAT), also known as sleeping sickness, is still a major concern in endemic countries. Its cyclical vector are biting insects of the genus *Glossina* or tsetse flies. In Guinea, the mangrove ecosystem contains the main HAT foci of Western Africa. There, the cyclical vector is *Glossina palpalis gambiensis*. A still ongoing vector control campaign (VCC) started in 2011 in the focus of Boffa, using tiny targets, with a 79% tsetse density reduction in 2016 and significant impact on the prevalence of the disease (from 0.3% in 2011 to 0.11% in 2013, 0.0352% in 2016 and 0.0097% in 2019). To assess the sustainability of these results, we have studied the impact of this VCC on the population biology of *G. p. gambiensis* in Boffa. We used the genotyping at 11 microsatellite markers and population genetic tools of tsetse flies from different sites and at different dates before and after the beginning of the VCC. In variance with a significant impact of VCC on the apparent densities of flies captured in the traps deployed, the global population of *G. p. gambiensis* displayed no variation of the sex-ratio, no genetic signature of control, and behaved as a very large population occupying the entire zone. This implies that targets deployment efficiently protected the human populations locally, but did not impact tsetse flies where targets cannot be deployed and where the main tsetse population exploits available resources. We thus recommend the pursuit of vector control measures with the same strategy, through the joint effect of VCC and medical surveys and treatments, in order to protect human populations from HAT infections until the disease can be considered as entirely eradicated from the focus.

## Introduction

Human African trypanosomosis (HAT), also known as sleeping sickness, is still a major concern for the WHO (Holmes, 2014; Simarro et al., 2015). Its agent is an euglenozoan kinetoplastid parasite, *Trypanosoma brucei gambiense* 1 (Jamonneau et al., 2019) (Tbg1), with a clonal propagation (Koffi et al., 2009, 2015), which recently emerged 10,000 years ago and spread from West Africa (Weir et al., 2016). It encompasses a complex life cycle, with a phase in blood sucking insects of the genus *Glossina* (Diptera, Hippoboscoidea), its cyclical vector (Bouyer et al., 2015).

In Guinea, the mangrove ecosystem still contains the main HAT foci of Western Africa (Simarro et al., 2015). There, *Glossina palpalis gambiensis* is the only known vector of the disease (Courtin et al., 2015). In Boffa, one of the three active HAT foci of the country (Kagbadouno et al., 2012), an active vector control campaign (VCC), aiming at reducing the human/tsetse contact, and subsequently at interrupting the transmission of Tbg1, was implemented in 2011 on the East bank of the Pongo river (Courtin et al., 2015). It is still ongoing and was extended to the whole focus in 2016 (Camara et al., 2021). This VCC was clearly successful, with a 80% reduction in individual catch per day, a 75% reduction of human exposure to tsetse bites, and a 70% reduction of the prevalence of the disease in the East bank of the Pongo river before 2016 (Courtin et al., 2015). Since 2016, for the whole focus, number of HAT cases dropped from 0.0352% to 0.0097% in 2019 (Camara et al., 2021). As regard to the vertebrate host, the exact role of animal or asymptomatic human reservoirs is still uncertain. Nevertheless, the genetic diversity of this parasite maintained by the Guinean populations suggests the existence of a significant amount of clonal lineages outside the patients involved in surveys (Koffi et al., 2009), in human and/or animal reservoirs. Several other studies appeared to confirm this hypothesis (Bucheton et al., 2002; Büscher et al., 2018).

A recent study of *G. palpalis palpalis* in the focus of Bonon (Côte d’Ivoire), revealed that a special allele at a given trinucleotide locus (GPCAG), experienced a brutal increase in frequency after control (Berté et al., 2019). This suggested the selection for resistance against control measure, associated with this particular allele.

Given these observations, deciding what strategies to use in the future will depend on the most precise knowledge we can get on the population impact of medical campaigns and VCC.

In this paper, we investigated the impact of VCC over 11 years on the HAT focus of Boffa (Figure 1), on capture densities and sex-ratio, as well as on *G. palpalis gambiensis* population genetics, in particular on locus GPCAG. We then discuss the best strategy to be used to optimize the protection of human populations from this neglected tropical disease.

**Figure 1:**
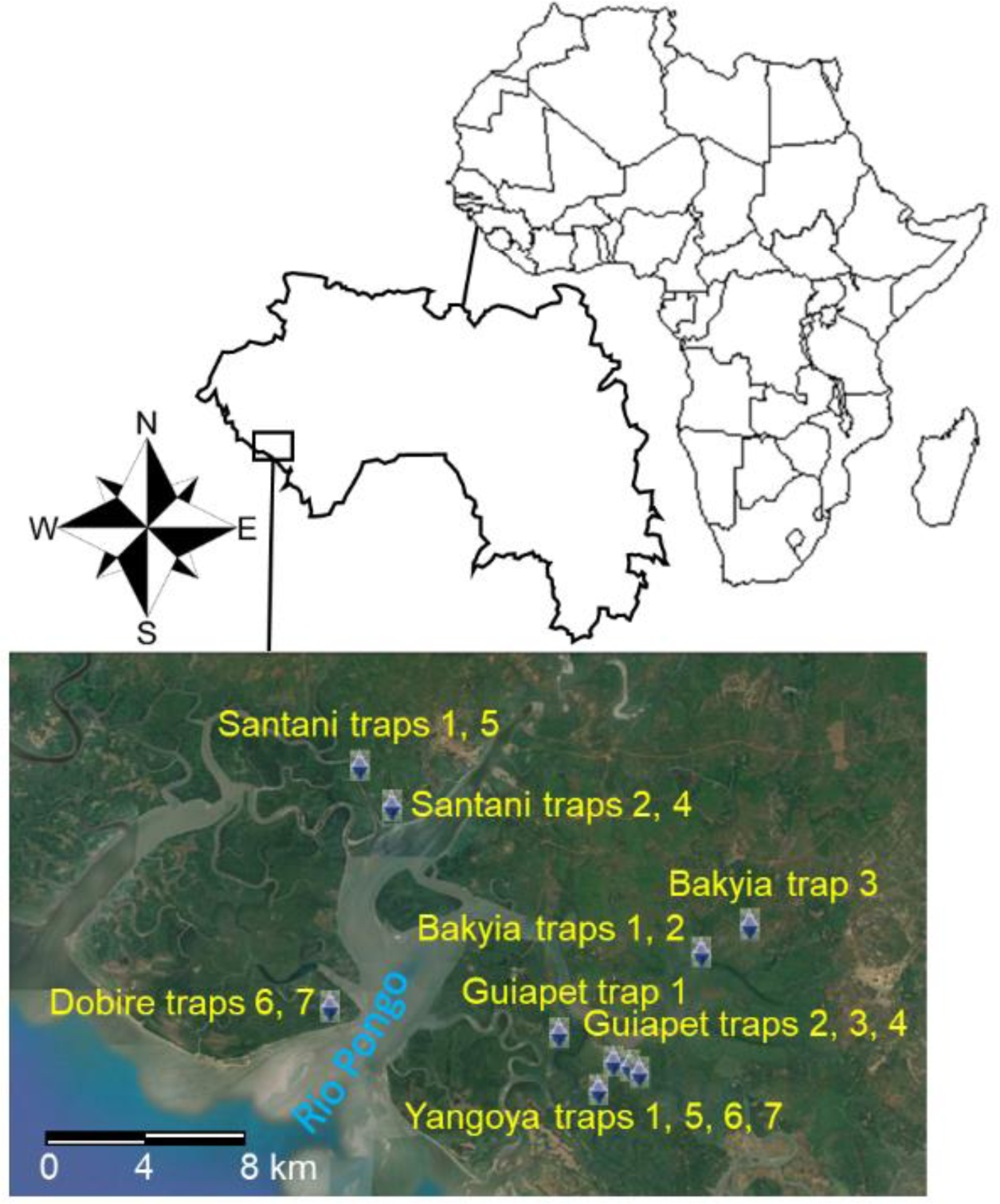
Localization of traps for the capture of *Glossina palpalis gambiensis* in the HAT focus of Boffa in Guinea. Dates of capture, number of flies genotyped, and other geographical information can be found in the Table 1.

**Table 1:** Number of genotyped *Glossina palpalis gambiensis* (*N*) in the different traps of different sites and date of capture in the HAT Focus of Boffa (Guinea). The cohort (considering six generations of tsetse flies per year) and the time (T) after the beginning of control in months (0=before), the bank of the Rio Pongo river (see Figure 1) and the land type are also indicated.

| Site | Trap | Date | Cohort | T | Bank | Land | N | N C-T-Bank |
| --- | --- | --- | --- | --- | --- | --- | --- | --- |
| Guiapet | 4 | 10/2009 | 0 | 0 | Left | Island | 23 | C0-T0-LeftBank |
| Yangoya | 1 | 10/2009 | 0 | 0 | Left | Island | 30 | 53 |
| Bakyia | 1 | 05/2011 | 10 | 0 | Left | Continent | 11 | C10-T0-LeftBank<br>65 |
| Bakyia | 2 | 05/2011 | 10 | 0 | Left | Continent | 8 |  |
| Bakyia | 3 | 05/2011 | 10 | 0 | Left | Continent | 5 |  |
| Guiapet | 2 | 05/2011 | 10 | 0 | Left | Island | 7 |  |
| Guiapet | 3 | 05/2011 | 10 | 0 | Left | Island | 4 |  |
| Yangoya | 1 | 05/2011 | 10 | 0 | Left | Island | 7 |  |
| Yangoya | 5 | 05/2011 | 10 | 0 | Left | Island | 3 |  |
| Yangoya | 6 | 05/2011 | 10 | 0 | Left | Island | 2 |  |
| Yangoya | 7 | 05/2011 | 10 | 0 | Left | Island | 16 |  |
| Yangoya | 8 | 05/2011 | 10 | 0 | Left | Island | 2 |  |
| Dobire | 6 | 05/2011 | 10 | 0 | Right | Island | 20 | C10-T0-RightBank<br>66 |
| Dobire | 7 | 05/2011 | 10 | 0 | Right | Island | 12 |  |
| Santani | 2 | 05/2011 | 10 | 0 | Right | Continent | 10 |  |
| Santani | 4 | 05/2011 | 10 | 0 | Right | Continent | 4 |  |
| Santani | 5 | 05/2011 | 10 | 0 | Right | Continent | 20 |  |
| Bakyia | 2 | 10/2019 | 60 | 92 | Left | Continent | 4 | C60-TX-LeftBank<br>35 |
| Guiapet | 2 | 10/2019 | 60 | 92 | Left | Island | 2 |  |
| Guiapet | 3 | 10/2019 | 60 | 92 | Left | Island | 11 |  |
| Yangoya | 1 | 11/2019 | 60 | 93 | Left | Island | 2 |  |
| Yangoya | 5 | 11/2019 | 60 | 93 | Left | Island | 4 |  |
| Yangoya | 6 | 11/2019 | 60 | 93 | Left | Island | 1 |  |
| Yangoya | 7 | 11/2019 | 60 | 93 | Left | Island | 11 |  |
| Dobire | 6 | 11/2019 | 60 | 33 | Right | Island | 9 | C60-TX-RightBank<br>27 |
| Dobire | 7 | 10/2019 | 60 | 32 | Right | Island | 1 |  |
| Santani | 2 | 11/2019 | 60 | 33 | Right | Continent | 8 |  |
| Santani | 4 | 11/2019 | 60 | 33 | Right | Continent | 1 |  |
| Santani | 5 | 11/2019 | 60 | 33 | Right | Continent | 8 |  |
| Bakyia | 2 | 10/2020 | 66 | 104 | Left | Continent | 3 | C66-TX-LeftBank<br>24 |
| Guiapet | 1 | 10/2020 | 66 | 104 | Left | Island | 15 |  |
| Guiapet | 2 | 10/2020 | 66 | 104 | Left | Island | 1 |  |
| Yangoya | 7 | 10/2020 | 66 | 104 | Left | Island | 5 |  |
| Dobire | 1 | 10/2020 | 66 | 44 | Right | Island | 6 | C66-TX-RightBank<br>20 |
| Dobire | 5 | 10/2020 | 66 | 44 | Right | Island | 2 |  |
| Santani | 1 | 10/2020 | 66 | 44 | Right | Continent | 10 |  |
| Santani | 3 | 10/2020 | 66 | 44 | Right | Continent | 2 |  |

N C-T-Bank: Names and numbers of riverbank subsamples, with the cohort (C) and the time (T) before (0) or after (X) the beginning of control.

## Material and Methods

### Data collection

All sites used for the entomological survey, number of captured flies and their gender can be seen in Kagbadouno et al (2012), Courtin et al. (2015), Camara et al. (2021), and, for most recent captures, in Supplementary file S1. All data and description of VCC in Boffa are available in two papers (Courtin et al., 2015; Camara et al., 2021) and the associated supplementary files of these articles.

Localization, date of sampling and number of genotyped flies can be found in Figure 1 and Table 1. A total of 290 flies were genotyped at 11 loci. Vector control was implemented with tiny targets from February 2012 in the sites of the left bank of Rio Pongo river and then on both sides after 2016. Then, according to Table 1, flies from 2009 and 2011 samples correspond to T0 (before control) on both sides of Rio Pongo river, while samples from 2019 and 2020 correspond to TX (after control has begun), on both sides of the river.

Active surveillance began in 2011, where 2730 individuals (1474 females and 1256 males) were captured. In 2019, 143 flies (80 females and 63 males) and in 2020, 215 (106 females and 109 males) were caught. The details on number of flies in the different dates and locations, and all genotypes, are available in the supplementary file S1.

Assuming a two month generation time (Williams, 1990), as can be seen from Table 1, the genotyped subsamples extended from cohort 0 (2009), 10 (2011), 60 (2019) and 66 (2020). Because individuals belonging to one of these cohorts have no chance of interacting with any individual from another of these cohorts, we then always considered these four cohorts as belonging to distinct subsamples.

Homogeneity of sex-ratio was tested across cohorts with a Fisher exact test with Rcmdr.

### Genotyping

Three legs of each tsetse fly were subjected to chelex treatment to obtain DNA for genotyping (Ravel et al., 2007). Genotyping was undertaken for 11 loci: X55-3 (Solano et al., 1997), XpGp13, pGp24 (Luna et al., 2001), GPCAG (Baker & Krafsur, 2001), pGp15, pGp16, and pGp27 (Ravel et al., 2020). Loci A10, B3, XB104, C102 were kindly provided by A. Robinson (Insect Pest Control Sub-program, Joint Food and Agriculture Organization of the United Nations/International Atomic Energy Agency Program of Nuclear Techniques in Food and Agriculture). Despite used at several occasions (Camara et al., 2006; Bouyer et al., 2007; Solano et al., 2009; Melachio et al., 2011), information about these loci were never published. Readers will find this information in Table A1 in the Appendix. Locus name with an X means that the locus is located on the X chromosome, and thus haploid in males. Cells with a “0” correspond to individuals with no amplification at the concerned locus, while uninterpretable profiles were coded “NA” (see Supplementary file S1).

### Data analyzed

Genotyping was undertaken in different ways and under different conditions for the different dates of sampling (see Table S1). For samples collected in 2009 and 2011, allele bands were resolved on a 4300 DNA Analysis System (LICOR, Lincoln, NE) using two infrared dyes (700 and 800nm) as described in (Solano et al., 2009). For samples collected in 2019 and 2020, an ABI 3500xl sequencer (Applied Biosystem, Waltham, Massachusetts) was used with four different dyes (FAM, NED, VIC and PET) as described in Berté et al. (2019). Consequently, the 11 loci were not analyzed with the same dyes between samples from 2009-2011 and samples from 2019-2020. Because it is generally admitted that the dye used can modify the apparent size of the amplified alleles, we genotyped again some samples from 2009-2011 at the different loci using the ABI 3500xl sequencer to recode all the genotypes from the samples from 2009-2011. In some subsets, too many missing data were observed at some loci or subsamples. Consequently, to avoid multiple analyses, and to keep a reasonable number of loci and for the sake of consistency, population genetic analyses were undertaken with females only, at the eight loci that were successfully amplified over all the dataset (see Table S1): X55-3, XpGp13, pGp24, A10, B3, XB104, C102, and GPCAG.

Unless specified otherwise, the global dataset (File S1) was processed with Create (Coombs et al., 2008) to produce datasets in different formats, depending on the kind of analyses to be undertaken.

### Selection of the relevant subsample units

The reproductive system of a population from a genotyped sample can be assessed through Wright’s *F*_IS_ (Wright, 1965), which is a measure of the relative inbreeding of individuals as compared to the subpopulation they belong to. This parameter was estimated with Weir and Cockerham’s unbiased estimator *f* (Weir & Cockerham, 1984), that we kept labelling *F*_IS_ for the sake of simplicity. In case of random mating in a dioecious population, a negative value is expected (Robertson, 1965; Pudovkin et al., 1996; Balloux, 2004; De Meeûs & Noûs, 2023). In tsetse flies, the trap can be the unit at which a significant subdivision may occur (Ravel et al., 2023), including *G. palpalis gambiensis* (Bouyer et al., 2009). Here, traps, and even sites, in combination with the cohort, often contained very few flies. To determine if traps, sites or river banks (right or left of the Rio Pongo River, see Figure 1) really mattered, we used the Wahlund effect approach (Goudet et al., 1994; De Meeûs et al., 2006). We compared the *F*_IS_T_ in traps, the *F*_IS_S_ in sites (ignoring traps), the *F*_IS_B_ within the two riverbanks (ignoring sites), and the *F*_IS_C_ within the four cohorts (0, 10, 61 and 67) (ignoring river banks). For this, we undertook a Friedman two-way analysis of variance by ranks for data paired by locus with the package R-commander (Fox, 2005, 2007) (rcmdr) for R (R-Core-Team, 2022). We then undertook planned one sided Wilcoxon signed rank test for paired data. If traps and/or sites and/or river banks matter, we expect a Wahlund effect (De Meeûs et al., 2007; De Meeûs, 2018) when pooling individuals from different origins (traps, and/or sites and/or river banks of the same cohort), meaning that the alternative hypothesis was *F*_IS_T_< *F*_IS_S_<*F*_IS_B_<*F*_IS_C_. When necessary (significant *p*-values), we corrected the *p*-values with the Benjamini and Yekutieli correction (Benjamini & Yekutieli, 2001) with R (command “p.adjust“), for test series with dependency.

### Quality testing of loci and subsamples

Quality testing of the subsamples and genotyping was assessed with linkage disequilibrium (LD), Wright’s *F*_IS_ and *F*_ST_ (Wright, 1965) and their variance across loci.

Linkage disequilibrium was tested with the *G*-based randomization test between each locus pair across subsamples as described in De Meeûs et al. (2009), with 10,000 random shuffling of genotypes of the two loci of each pair. Because there are as many non-independent tests as locus pairs, we adjusted the *p*-values with the Benjamini and Yekutieli (BY) procedure with R (command “p.adjust“).

Wright *F*-statistics were estimated with Weir and Cockerham’s unbiased estimators (Weir & Cockerham, 1984): *f* for *F*_IS_, *θ* for *F*_ST_ and *F* for *F*_IT_. Their significant deviation from the expected value under the null hypothesis (H0: *F*_IS_=0 or *F*_ST_=0) was tested with 10,000 randomizations of alleles between individuals within subsamples (for *F*_IS_) and of individuals between subsamples (for *F*_ST_). For *F*_IS_, the statistic used was Weir and Cockerham’s *f* for each locus and overall. For testing subdivision we used the *G* based test (Goudet et al., 1996) for each locus and overall. For *F*_IS_, tests were one-sided (*F*_IS_>0), and two-sided *p*- values were obtained by doubling the one-sided *p*-value if <0.5 or we doubled 1-*p*-value otherwise. Confidence intervals (95%CI) were computed with 5000 bootstraps over loci for the average over loci, and over individuals for the confidence intervals around each locus. The standard error of jackknives over loci (SE) was computed for *F*_IS_ and *F*_ST_ (SE_*F*_IS_ and SE_*F*_ST_ respectively) for null allele diagnostics (see below).

All this computations and randomizations were undertaken with Fstat 2.9.4 (Goudet, 2003), updated from Fstat 1.2 (Goudet, 1995), except the bootstraps over individuals, for which we used Genetix (Belkhir et al., 2004). Since Genetix computes bootstraps for each locus in each subsamples, we averaged these values across subsamples to obtain 95%CI of *F*_IS_ for each locus.

Null allele signatures were looked for with several criterions: the ratio of SE of *F*_IS_ over *F*_ST_ (*r*_SE_), as computed with jackknives over loci; the correlation between *F*_IS_ and *F*_ST_, and between *F*_IS_ and the number of missing genotypes (*N*_b_) (De Meeûs, 2018). These correlations were tested with a one-sided (positive correlation) Spearman’s signed rank correlation test. The regression *F*_IS_∼*N*_b_ was also undertaken to determine the proportion of the variance of *F*_IS_ that was explained by missing genotypes (putative null homozygotes) with the determination coefficient *R*², and to extrapolate the possible basic *F*_IS_ (*F*_IS_0_) in absence of null alleles through the intercept of this regression. Null allele frequencies were also estimated with the EM algorithm (Dempster et al., 1977) implemented by FreeNA (Chapuis & Estoup, 2007). For these analyses, missing genotypes (true blanks) were coded as null homozygotes for allele 999, as recommended (Chapuis & Estoup, 2007). The goodness of fit of observed blank genotypes, with the expected number of missing data, was assessed with a one-sided exact binomial test with R (command “binom.test“). If null alleles and random mating fully explain the data, there should be at least as many blank genotypes as expected. For this test we computed the square of null allele frequency of each locus in each subsample and multiplied it by the subsample size. To increase statistical power, we summed these expected number of null homozygotes over all subsamples (*N*_Exp-b_) and compared those to the observed number of missing genotypes for each locus, over all the considered dataset. We then also undertook the regression *F*_IS_∼*p*_n_, where *p*_n_ is the global null allele frequency.

Stuttering detection and correction followed the procedure described elsewhere (De Meeûs et al., 2021; De Meeûs & Noûs, 2022) over all subsamples and for each locus, using the associated template “TestStutterDioecious-n1000N100-1-10%Stuttering.xlsx” available at https://zenodo.org/record/7029324, exact binomial tests and adjustment of *p*- values with Benjamini and Hochberg (BH) procedure with R (command “p.adjust“).

Short allele dominance (SAD) was looked for with the correlation method between *F*_IT_ and the size of the alleles (Manangwa et al., 2019) using a one-sided (negative relationship) Spearmans rank correlation test. In case of doubt, we also undertook the regression *F*_IS_∼Allele size, weighted with *p_i_*(1-*p_i_*), and where *p_i_* was the frequency of the allele (De Meeûs et al., 2004). Correlation and regression tests were undertaken with rcmdr (Fox, 2005, 2007). For SAD, the regression allows to minimize spurious correlations due to rare alleles. In rcmdr, regression tests are two-sided. To obtain a one-sided *p*-value, when the slope was negative, we simply halved the two-sided *p*-value, or computed 1-*p*- value/2 otherwise. We labelled the correlation *p*-value *p*_cor_ and the regression one *p*_reg_.

### Subdivision

Comparisons of *F*-statistics between different subsamples or between *F*_IS_ and *F*_IT_ was undertaken with Wilcoxon signed rank tests for paired data with rcmdr, using the loci as pairing factor. Tests were one-sided in the appropriate direction (e.g. *F*_IT_>*F*_IS_). Subdivision was measured and tested with Wright’s *F*_ST_ and the *G* randomization test, as described above. Genetic differentiation was measured and tested between each pair of cohorts in the same way, except that systematic paired test required BY corrections for significant *p*-values.

### Effective population sizes and effective population densities

We estimated effective population sizes (*N_e_*) with several algorithms and softwares. The first method is the heterozygote excess method from De Meeûs and Noûs (De Meeûs & Noûs, 2023), computed for each locus in each cohort, ignoring values below or equal to 0, and averaged across loci, for each subsample (e.g. cohort). The second method was the linkage disequilibrium based method (Waples & Do, 2010), corrected for missing data (Peel et al., 2013), implemented by NeEstimator (Do et al., 2014), and assuming random mating and selecting results for alleles with frequency above or equal to 5%. The third method was the co-ancestry method (Nomura, 2008), also implemented by NeEstimator. The fourth method used the averages obtained with four temporal methods. Three were implemented in NeEstimator: Pollak, Nei and Tajima, and Jorde and Ryman (Nei & Tajima, 1981; Pollak, 1983; Jorde & Ryman, 2007) assuming random mating and selecting results for alleles with frequency above or equal to 5%. For these three temporal method, we averaged *N_e_* across the different comparisons (different cohort pairs), ignoring “Infinite” outputs. The last temporal method was the maximum likelihood temporal method implemented in MLNe (Wang & Whitlock, 2003). For MLNe, maximum and minimum values corresponded to the confidence interval outputted by the procedure. We then computed the average across all temporal methods, weighted by the number of usable values and averaged minimum and maximum values in the same way. For MLNe, the weight was the number of cohort pairs (6). The fifth method was the within and between loci identity probabilities (Vitalis & Couvet, 2001a) with Estim (Vitalis & Couvet, 2001b). The sixth method was the sibship frequency method (Wang, 2009) with Colony (Jones & Wang, 2010), assuming female and male polygamy, and inbreeding. For each method, we computed the average, the minimum and maximum across usable values, and assigned a weight (number of usable values). For temporal methods, the weight was set to 4 (number of cohorts). We then obtained the grand average for *N_e_* and minimum and maximum (minimax) values through an average across methods, weighted by the aforementioned weights. This methodology obviously provided a rough approximation of *N_e_* and the space of its possible values, and finding the most accurate way will need specific multi-scenarios simulation approaches, as discussed elsewhere (De Meeûs & Noûs, 2023).

To compute effective population densities, we considered three different surfaces, computed with the function “insert a polygon” of Google Earth Pro. The smallest area was defined by the traps containing flies that were indeed genotyped, as in Figure 1 and Table 1, and labelled *S*_Genet_=132 km². The second surface was defined by all traps that captured at least one tsetse fly in 2011, during the most extensive survey campaign (Kagbadouno et al., 2012) (see File S1), *S*_C_=224 km². The third area was defined by the limits of the survey defined before 2011 (see File S1); *S*_L_=629 km². The largest surface was drawn considering all “mangrove-like” environments in Google Earth Pro, *S*_max_=1301 km². Effective population densities were then computed by dividing the effective population size by these surfaces. For the sake of comparison, we also computed the densities of captured flies in 2011 (initial survey, T0), in 20019 and 2020 (see Files S1 for raw data). We also compute the sex-ratios of captured (c) flies, *SR*=*N*_c-males_/*N*_c-females_ with these data. The homogeneity of the proportion of females across cohorts was tested with a Fisher exact test with rcmdr. The evenness of sex-ratios was then tested with a two-sided exact binomial test with R (command “binom.test“).

Finally, maximum distances between the most remote sites defined by each aforementioned surface was computed with Google Earth Pro with the menu “Add a trajectory“.

### Bottleneck signatures

We tried to find bottleneck signatures with the diversity excess method and the Wilcoxon signed rank test method with Bottleneck (Cornuet & Luikart, 1996), with a particular interest to subsamples genotyped after the beginning of control (cohorts 61 and 67). We used the three mutation models: IAM, TPM with default options, and SMM. Significant signatures of a bottleneck can be suspected if the test is highly significant with the IAM and significant with the TPM at least. Weaker signals tend to be produced in population with very small effective population sizes (De Meeûs, unpublished simulation results).

## Results

### Captured flies, apparent densities and sex-ratio

The number of flies captured, per gender and total, the apparent density per trap and day (ADTD) and the sex-ratio for the different cohorts are presented in Table 2. There is a clear drop in ADTP after the beginning of VCC. We also observed a significant change in sex-ratio across cohorts (*p*-value=0.0015), but this is only due to the small sex-ratio observed in 2009 (Table 2). When we removed 2009 data, differences are not significant any more (*P*-value=0.491).

**Table 2:** Number of males (*N*_M_), females (*N*_F_) and total number (*N*) of *Glossina palpalis gambiensis* captured in the HAT focus of Boffa (Guinea), with year of capture and corresponding cohort, apparent density per trap and day (ADTD) and sex-ratio.

| Year | Cohort | $N_M$ | $N_F$ | $N$ | ADTD | Sex-ratio |
| --- | --- | --- | --- | --- | --- | --- |
| 2009 | 0 | 41 | 97 | 138 | 23.00 | 0.42 |
| 2011 | 10 | 1246 | 1484 | 2730 | 25.87 | 0.84 |
| 2019 | 60 | 62 | 82 | 144 | 4.44 | 0.76 |
| 2020 | 66 | 68 | 68 | 136 | 3.58 | 1 |

### Defining the relevant geographic scales for subsample units

There was no significant difference between *F*_IS_T_, *F*_IS_S_ and *F*_IS_B_ (all *p*-values>0.4) (Figure 2). We thus ignored traps, sites and river banks for further analyses, and kept only the different cohorts (cohorts 0, 10, 60 and 66) as subsample units.

**Figure 2:**
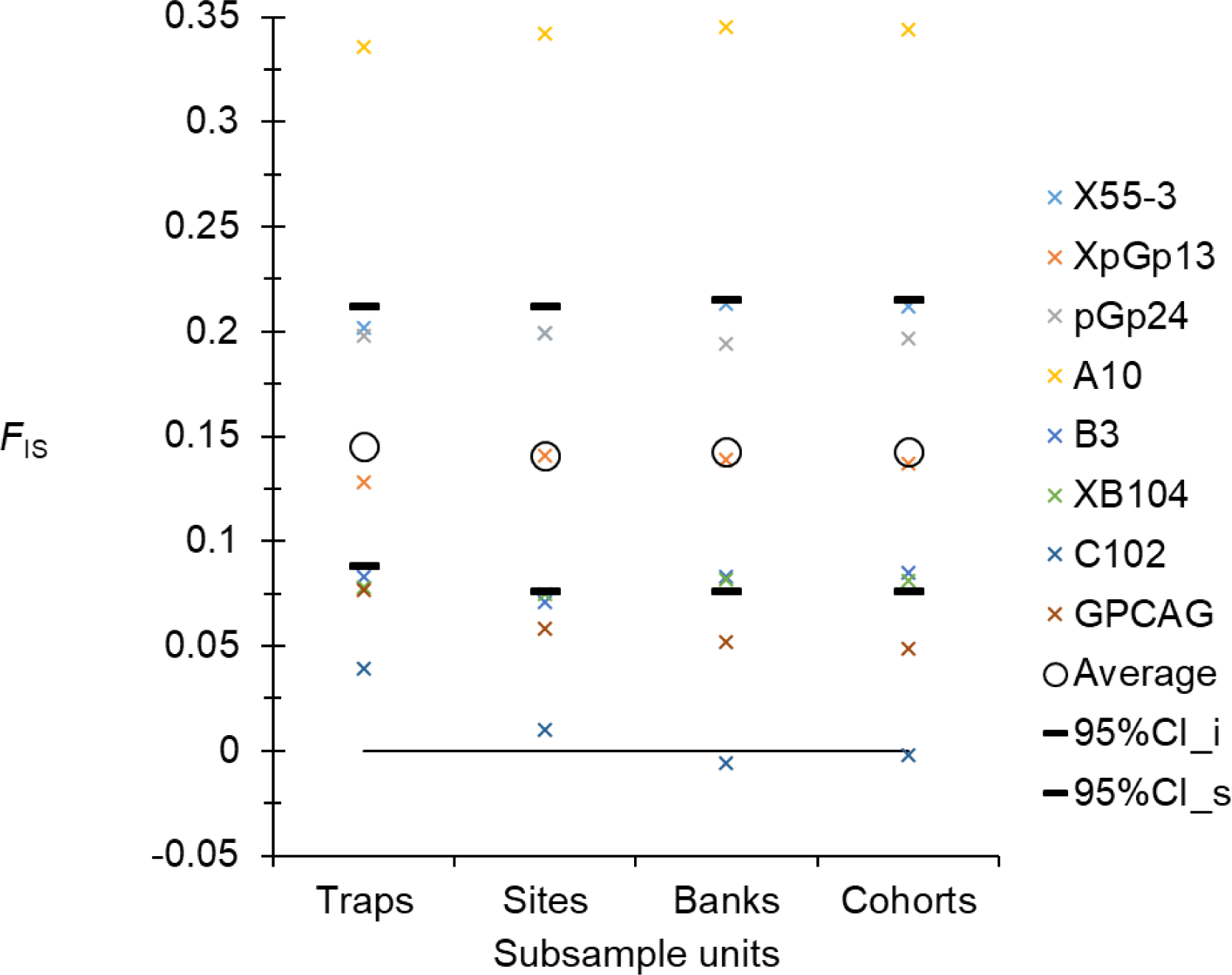
Comparisons of *F*_IS_ within traps (*F*_IS_T_), within sites (ignoring traps) (*F*_IS_S_), within river banks (ignoring sites) (*F*_IS_B_), and within each cohort (ignoring river banks) (*F*_IS_C_) in female subsamples of *Glossina palpalis gambiensis* across 11 years in the mangroves of Boffa, Guinea, for eight loci (crosses of different colors), average (empty circles) and 95%CI (black dashes). The global comparison outputted a *p*-value=0.9663 and none of paired tests provided a *p*-value<0.4.

### Quality testing of the subsamples and loci

Only one locus pair out of 28 appeared in significant LD (less than 4%) (*p*- value=0.039). It did not remain significant after BY correction. We thus considered all loci to be statistically independent from each other’s.

There was a substantial but variable heterozygote deficit: *F*_IS_= 0.143 in 95%CI=[0.076, 0.215] (*p*-value<0.0002) (Figure 3). With *r*_SE_=19, and the positive correlation between *F*_IS_ and *F*_ST_ (*ρ*_=_0.5988, *p*-value=0.0584), we could suspect that most of this heterozygote deficit could be explained by null alleles. Nevertheless, the correlation between *F*_IS_ and *N*_b_ did not seem to confirm this (*ρ*=0.0952, *p*-value=0.4201). Nonetheless, this lack of signal resulted from three outlier loci (pGp24, B3 and C102), which displayed too many missing genotypes as regard to their relatively low *F*_IS_ (Figure 3). This means that many of these missing data, at these three loci, probably did not correspond to true null homozygotes.

**Figure 3:**
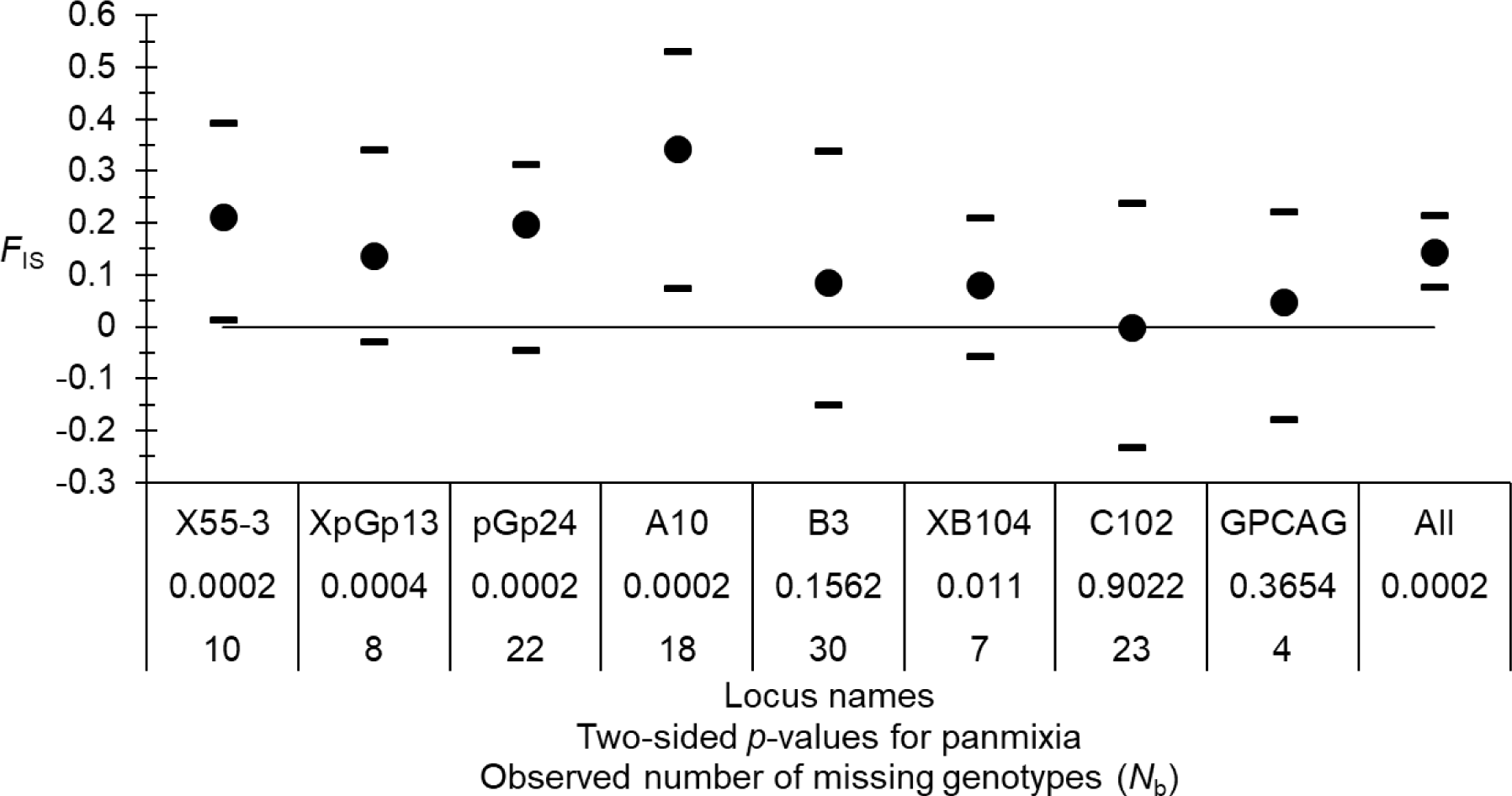
Variation of *F*_IS_ across loci in the four cohorts of *Glossina palpalis gambiensis* from Boffa (Guinea). Averages (black dots), and 95% confidence intervals (black dashes) were computed by 5000 bootstraps over individuals (for each locus) and over loci (All). The two sided *p*-values for significant deviation from panmixia and the number of observed missing genotypes are also provided.

We thus undertook the correlation and the regression *F*_IS_∼*N*_b_ without these three loci. The results can be observed in the Figure 4. The correlation became significant and the regression suggested that missing genotypes (null alleles) explains almost all (96%) of *F*_IS_ variation. The intercept was negative, as expected in small, pangamic dioecious populations (*F*_IS_0_=-0.0412 in 95%CI=[-0.1872, 0.1171]).

**Figure 4:**
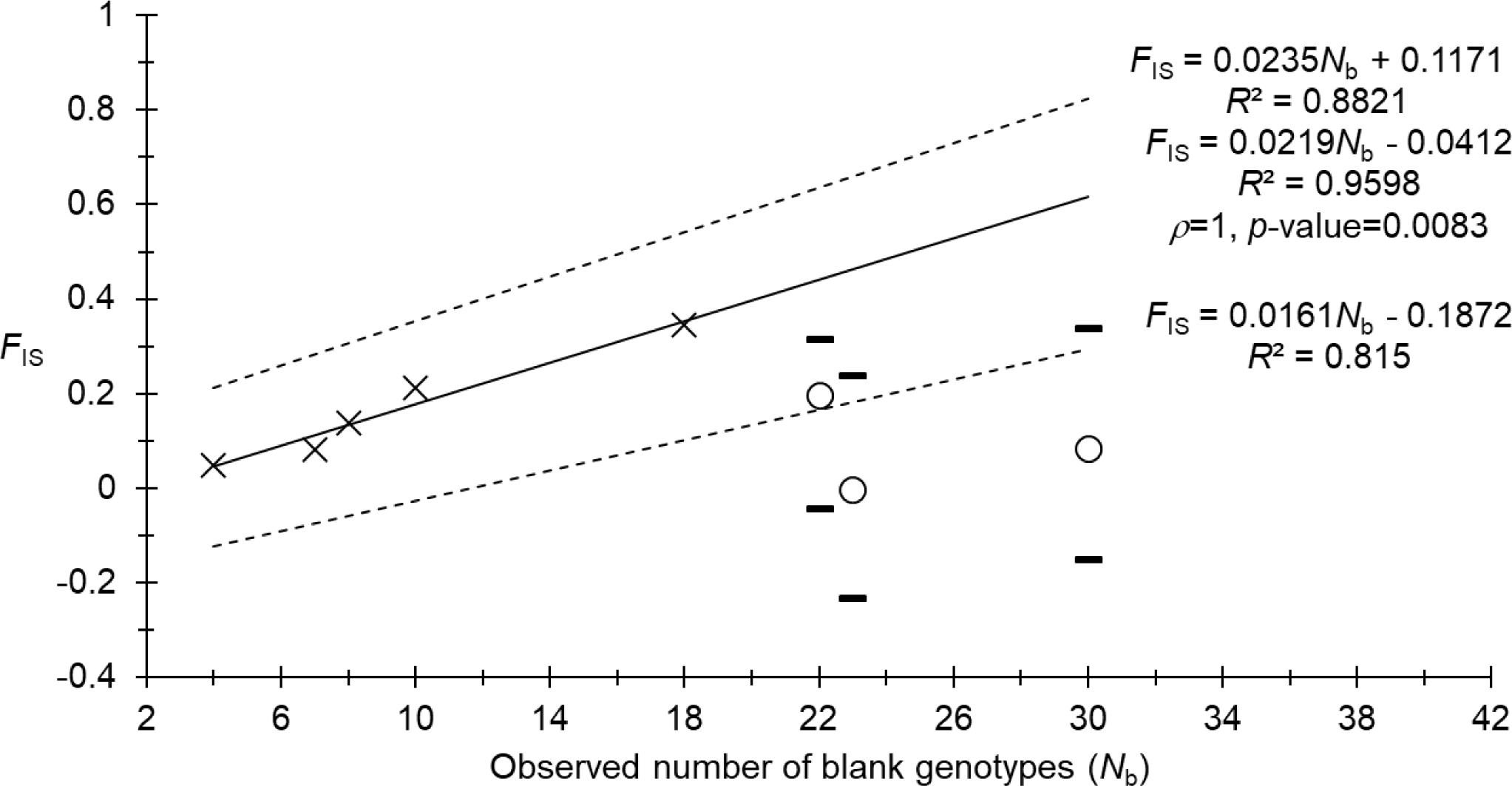
Regression of *F*_IS_ by the number of missing genotypes (black crosses and continuous line) without loci displaying an excess of missing genotypes (pGp24, B3 and C102) (empty circles) and of the corresponding 95%CI of bootstraps over individuals for *Glossina palpalis gambiensis* from Boffa (Guinea) (black dashes and broken lines).

For null allele frequency estimates, we recoded as homozygous for the null allele (allele coded 999) all missing genotypes but for the three outlier loci. Indeed, for these loci, most missing data (coded “0“, and kept as such) probably corresponded to another problem. With these data, estimate of null allele frequencies, and the resulting expected number of missing data over all subsamples fitted very well with the observed ones at each locus (all *p*-values>0.2). Moreover, the regression *F*_IS_*∼p*_n_ displayed very good results (*R*²=0.8566, *F*_IS_0_=0.012 in 95%CI=[-0.1951, 0.2143]). In this regression, loci pGp24, B3 and C102 were not outliers anymore. Nevertheless, the impossibility to determine which missing data are true null homozygotes at these loci provided a drop in accuracy. Without those, the regression was improved (*R*²=0.9851, *F*_IS_0_=-0.0391 in 95%CI=[-0.1891, 0.1151]).

We also undertook the stuttering and SAD analyses. We found a significant stuttering signature for locus A10 only (*p*-value=0.0286). Nevertheless, null alleles explained this locus well enough (Figures 3 and 4) so that we could consider this result as coincidental. Three loci displayed marginally significant or not significant signatures of SAD: X55-3 (*p*_cor_=0.0423, *p*_reg_=0.1551), pGp24 (*p*_cor_=0.0963, *p*_reg_=0.0489), and B3 (*p*_cor_=0.288, *p*_reg_=0.4841). Given the weakness of such results, and that these three loci were very well explained by null alleles, we did not consider that any locus was affected by SAD, and that a few null alleles randomly associated with small alleles provided these weak signatures.

We detected no Wahlund effect and the significant heterozygote deficits observed were entirely caused by null alleles of frequencies 0.02 to 0.28.

Average subdivision was globally not significant with a weak variance across the different loci: *F*_ST_=-0.001 in 95%CI=[-0.004, 0.004] (*p*-value=0.6938). In particular, GPCAG displayed a negative value: *F*_ST_=-0.007 (-0.006 with FreeNA correction) (*p*-value=0.1709), and allele 219 (here labelled 220, due to probable lag produced by the change in size calibration on this species), did not increase in frequency after control had begun: 0.094, 0.093, 0.065, and 0.022 for cohorts 0, 10, 60 and 66, respectively.

### Subdivision

As demonstrated above, we could not detect any geographic subdivision signature. Subdivision was globally weak across the different cohorts: average *F*_ST_=-0.001 in 95%CI=[-0.004, 0.004] and not significant (*p*-value=0.6938). Moreover, *F*_IT_=0.142 in 95%CI=[0.073, 0.218] was not significantly bigger than *F*_IS_=0.143 in 95%CI=[0.076, 0.215] (*p*-value=0.7695).

We also measured and tested subdivision between each pair of cohorts. In the Figure 5, we can confirm the absence of any subdivision. Moreover, subdivision between cohorts before control (T0) and after control had begun (TX) did not appear bigger than for other cohort comparisons.

**Figure 5:**
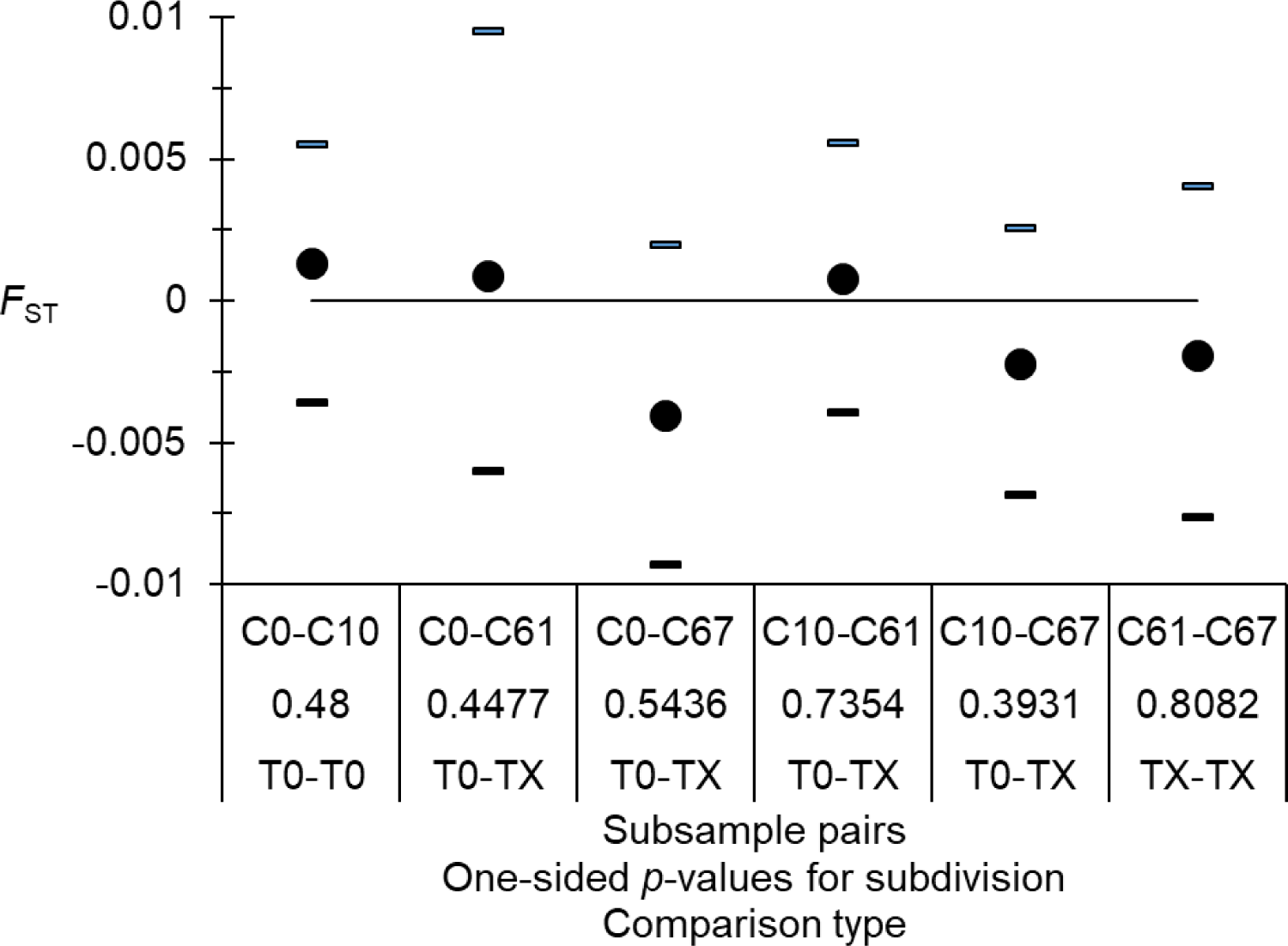
Subdivision measures (*F*_ST_) and test between each pair of cohorts (C0, C10, C60 and C66). The result of the test (*p*-values) and the comparison type between cohorts before control (T0) or after the beginning of control (TX) are also indicated.

### Effective population sizes and effective population densities

For this section, Estim provided no usable values. For the remaining methods, results are presented in the Table 3 and Figure 6. These values were quite variable, but did not display particularly smaller values after control (26 and 18 for cohorts 60 and 66 respectively) than before control (22 and 42 for cohorts 0 and 10 respectively). Nevertheless, such comparisons could only be made for the heterozygote excess, co-ancestries and sibships methods. The apparent increase in 2011 (cohort 10, before the VCC) can only be observed for three methods (heterozygote excess, Coancestries and Sibship). It is probably more due to sampling variances than to a real and brutal increase of the tsetse population.

**Figure 6:**
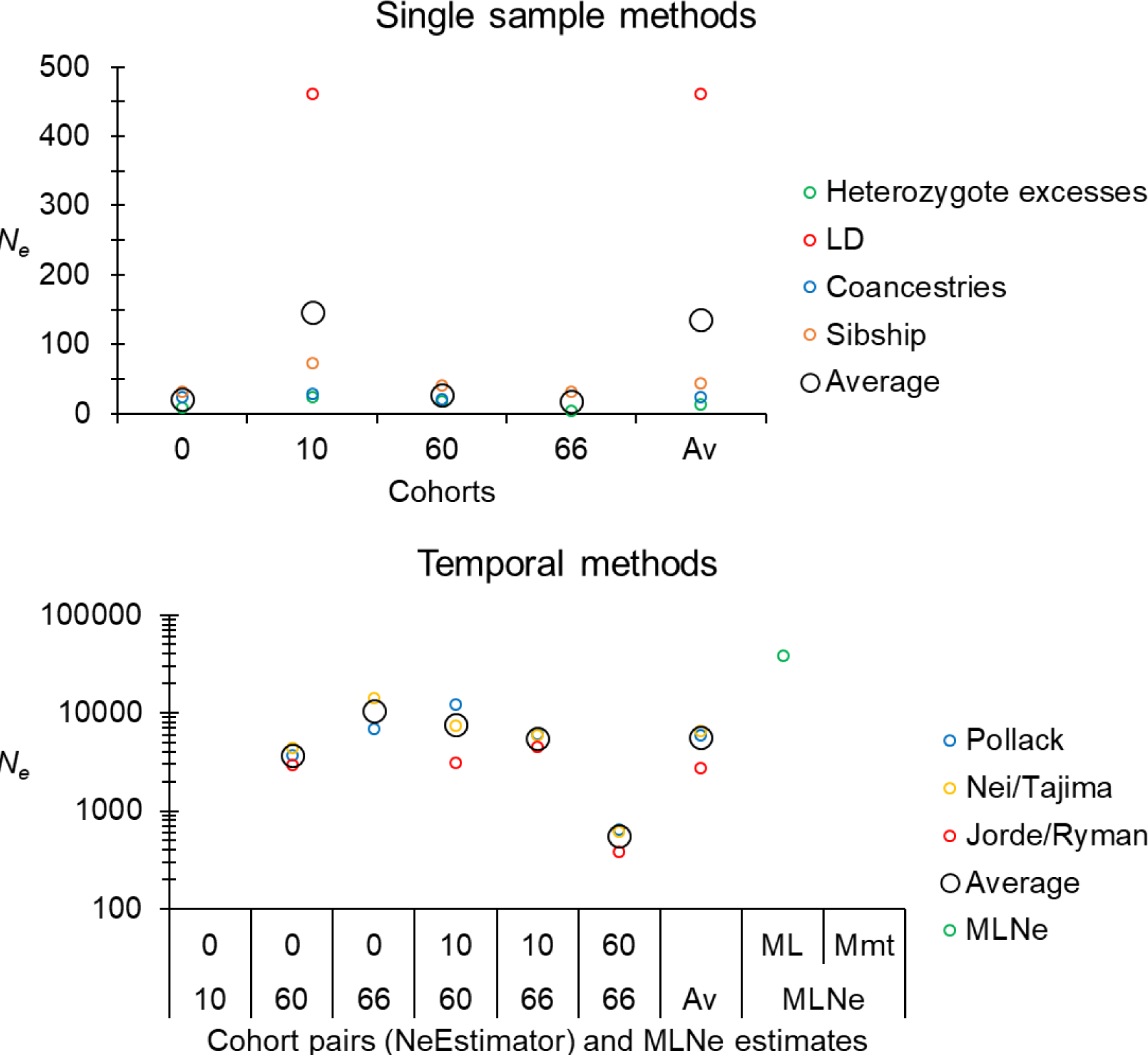
Comparisons of effective population sizes obtained with the different single sample and temporal samples methods for *Glossina palpalis gambiensis* from the sleeping sickness focus of Boffa, for the different cohorts and cohort pairs and unweighted averages (Av) over those, and for temporal methods implemented in MLNe, over all cohorts. Empty data means “Infinite” result. ML: maximum likelihood; Mmt: moment.

**Table 3:**
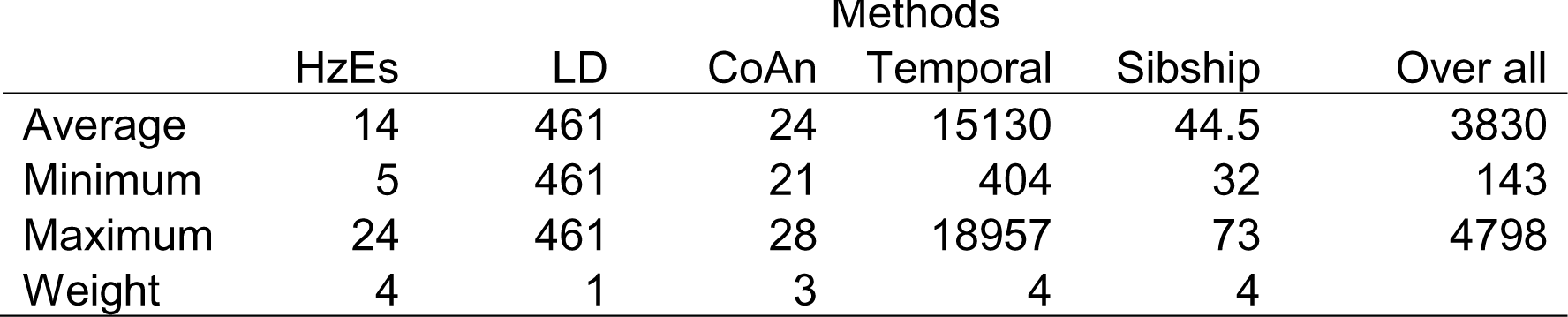
Effective population size, minimum and maximum values estimated with different methods, and over all method (average weighted with “Weight“). Codes for methods are HzEs: heterozygote excess; LD: linkage disequilibrium; CoAn: coancestries.

|  | Methods |  |  |  |  |  |
| --- | --- | --- | --- | --- | --- | --- |
|  | HzEs | LD | CoAn | Temporal | Sibship | Over all |
| Average | 14 | 461 | 24 | 15130 | 44.5 | 3830 |
| Minimum | 5 | 461 | 21 | 404 | 32 | 143 |
| Maximum | 24 | 461 | 28 | 18957 | 73 | 4798 |
| Weight | 4 | 1 | 3 | 4 | 4 |  |

Taking the surface occupied by genotyped flies, the resulting effective population density was *D_e_*_-Genet_=8 in minimax=[1, 17] flies per km². However, given the absence of any population subdivision signature, we considered that this quantity was a very strong overestimate. In the Figure 7, we can see the effective population densities computed with *S*_C_, *S*_L_ and *S*_max_. Such densities became much smaller when temporal methods were ignored (*D_e_*-_max_=0.05km² in minimax=[0.04, 0.06]).

**Figure 7:**
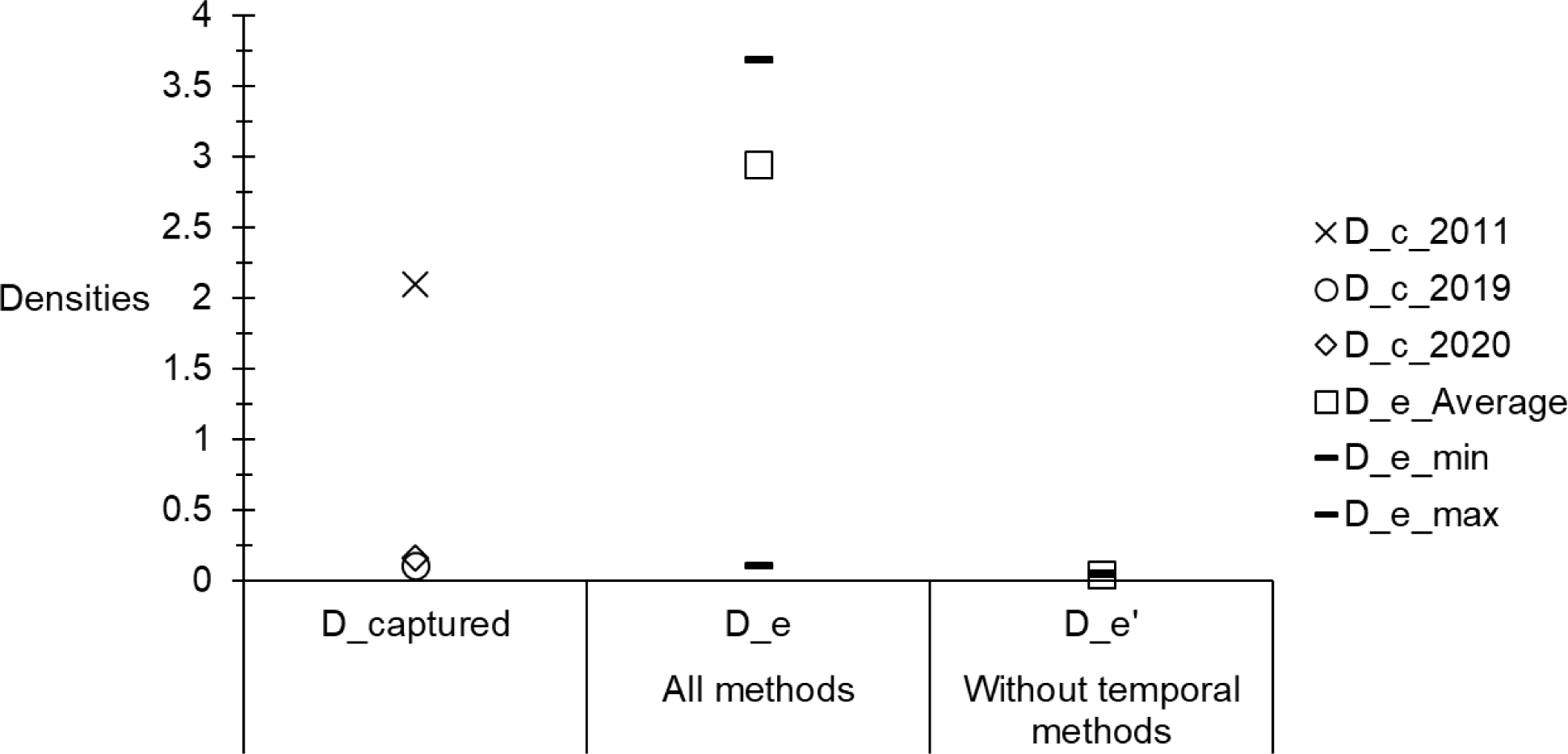
Densities of captured flies (Dc) and corresponding effective population densities (De) while considering the whole surface of Boffa (as defined in the text) occupied by tsetse flies in the HAT focus of Boffa (Guinea), for different years and different effective population size averages: with all methods, or without temporal methods.

There was no significant variations of the sex ratio across years (0.85, 0.79 and 1.03 for 2011, 2019 and 2020 respectively) (*p*-value=0.3253). Combining all years, captured flies displayed a highly significant female biased sex ratio *SR*=0.86 (*p*- value<0.0001).

Capturing flies after control had begun, in the same trapping sites, obviously appeared more difficult than before control (Figure 7).

The maximum distances between the most remote points were 19 km, 40 km, 44 km and 62 km for the areas *S*_Genet_, *S*_C_, *S*_L_ and *S*_max_ respectively. Depending on the true area occupied by the population of tsetse flies in Boffa, these may represent the average dispersal distances from one generation to the next.

### Bottleneck signatures

The results of these analyses are given in Table 4, where an absence of any bottleneck signature can be noticed, in particular after the beginning of control (cohorts 60 and 66).

**Table 4:**
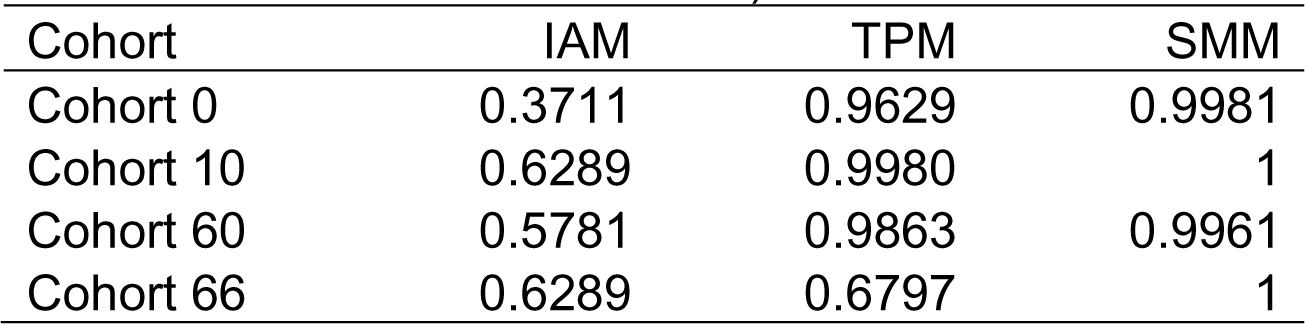
One sided *p*-values for the detection of a genetic signature of a bottleneck with the three models of mutation (IAM, TPM and SMM) for the different subsamples before the beginning of vector control (Cohorts 0 and 10) and after the beginning of control (Cohorts 60 and 66).

## Discussion

All heterozygote deficits observed at the eight loci that passed the quality control tests were entirely explained by null alleles. Wahlund effect or inbred systems of mating were dismissed by different observations: absence of linkage disequilibrium; variance of *F*_IS_ across loci, with several ones not significantly deviating from the panmictic model; negative intercept of the regression *F*_IS_*∼N*_b_. The eight loci studied here in female *G. palpalis gambiensis* converged with the existence of a large and stable pangamic population of this species occupying the HAT focus of Boffa. This was confirmed by the absence of any Wahlund effect when pooling all individuals from all the sampling area into one subsample in each cohort. The regression of the *F*_IS_ with null alleles or number of missing data, with a determination coefficient close to unity and a negative intercept confirmed this perception. No notable difference in effective population sizes from one cohort to the other could be observed. The apparently increase in cohort 10, before the beginning of the VCC, was probably due to sampling variance. Indeed, it is quite unlikely that, as brutal it could have been, an increase in the size of the population would present a significant signature in its effective population size after only 10 generations. If so, it is then hard to understand why no signature of any bottleneck was observed at generation 66. We could not observe any genetic differentiation between the different cohorts, whether within or between dates before or after the VCC had begun. This was in line with the extreme effective population size estimates with temporal methods, as compared to single sample based methods. The reality probably lies in between, which means that some genetic drift should occur, leading to some genetic differentiation. This may require more autosomal markers to be confirmed.

Subdivision tests also confirmed a free migration of tsetse flies in the area defined by this tsetse population. Taking the total surface where flies could be captured and the total surface of the survey, as the most realistic values, the corresponding dispersal distance would be between 40 and 44 km per generation. The constant shading, high humidity and wind conditions of the particular continuously favorable mangrove ecosystem probably explain such important values (Courtin et al., 2010, 2015; Courtin & Kagbadouno, 2011). Within these surfaces, the effective population density could be considered to vary between 3 and 6 individuals per km², in minimax≈[0.1, 8]. These range within the smallest values as compared to what can be found elsewhere (De Meeûs et al., 2019). This is even worse if we consider that, here, we could use temporal methods for estimating effective population sizes. Temporal methods are almost never available in the literature, but provided very high values in the present study. Without temporal methods, densities would have fallen to very small values: between 0.05 and 0.1 individuals per km² only, for the two surfaces respectively.

We observed a significant female biased *SR* in this population with small effective population densities. In a recent study of *G. fuscipes fuscipes* from Chad (Ravel et al., 2023), authors found a negative correlation between *SR* and densities. Scattered distribution of hosts, leading to smaller effective population densities of the flies, would impose to these insects to travel more to find blood meals. Because of larval feeding, female tsetse would need to feed more often than males and be captured more often than those in such kind of ecological frameworks. On the contrary, abundance of hosts would maintain dense populations of tsetse flies, with no need to spend time seeking for blood meals, especially so for females, while males still need looking for mates.

We observed the absence of any significant genetic differentiation between subsamples before and after the beginning of control, especially at locus GPCAG. Additionally, we failed to evidence any strong (if any) effect on effective population sizes before and after the beginning of VCC, and observed a total absence of any genetic signature of a bottleneck, even after a 79% drop in captured flies (Camara et al., 2021). All these results testify of an absence of consequences of the VCC on the genetic structure of this tsetse population, at the scale of the focus. Nevertheless, the dramatic drop in captured flies after the VCC had begun, indicated an efficient protection of the human population against tsetse bites around the targeted zones. The drop in tsetse bites and HAT prevalence and incidence in selected controlled zones has already proved such a protection (Courtin et al., 2015). It also indicates that in the areas targeted by vector control, tsetse densities would probably regain initial levels rapidly in case of scaling back. The fact that a significant decrease in HAT prevalence was observed (Courtin et al., 2015; Camara et al., 2021) also advocates for the benefic effects of the VCC. Finally, the absence of impact on the targeted population at the scale of the whole mangrove of Boffa, suggests that VCC as it was undertaken, is efficient at protecting human populations, but does not significantly affect the genetic diversity of this particular system. “Emptied” spots simply became reinvaded by a representative sample of the whole population, which may come from any part of the focus, even locally. Indeed, in the absence of any spatio-temporal population structure, it is impossible to determine where tsetse flies captured in one particular trap came from.

In conclusion, due to the difficulties to access to all tsetse-infested zones, where multiple hosts are available for these blood feeding insects, maintaining control measures in zones frequented by human populations seems a reasonable measure, in order to protect these populations from tsetse bites and subsequent transmission of HAT, as we know that these measures are indeed very protective (Courtin et al., 2015). Given that the existence of hidden human and/or animal reservoirs can be suspected (Koffi et al., 2009; Kaboré et al., 2011; Jamonneau et al., 2012; Büscher et al., 2018), and given that we have hereby shown that the HAT focus of Boffa harbors a freely circulating tsetse population, such a measure, together with medical surveys and treatment of infected patients, appears mandatory for now and a close future, before more research results shed more light on the eco-epidemiology of this deadly disease in mangrove ecosystems.

## Acknowledgements

The authors would like to thank Fabien Halkett and two anonymous referees for their comments that helped improve our manuscript.

## Author contributions

Moise S. Kagbadouno: data collection, data analyses and manuscript correction. Modou Séré: genotyping, data analyses and manuscript correction. Adeline Ségard: genotyping and manuscript correction. Abdoulaye Dansy Camara: data collection and manuscript correction. Mamadou Camara: supervision and manuscript correction. Bruno Bucheton: supervision, data collection and manuscript correction. Jean-Mathieu Bart: supervision, data collection and manuscript correction. Fabrice Courtin: supervision, data collection and manuscript correction. Thierry de Meeûs: supervision, data analyses, writing of the manuscript and design of figures. Sophie Ravel: supervision, genotyping and manuscript correction.

## Data, scripts, code, and supplementary information availability

Raw data are available at https://zenodo.org/record/8181166. For data analyses, all R scripts used are inserted in the main text. For other analyses we used facilities without the need of scripts (e.g. click and point programs or packages).

## Appendix

### Appendix 1 Information on unpublished microsatellite loci

**Table A1:**
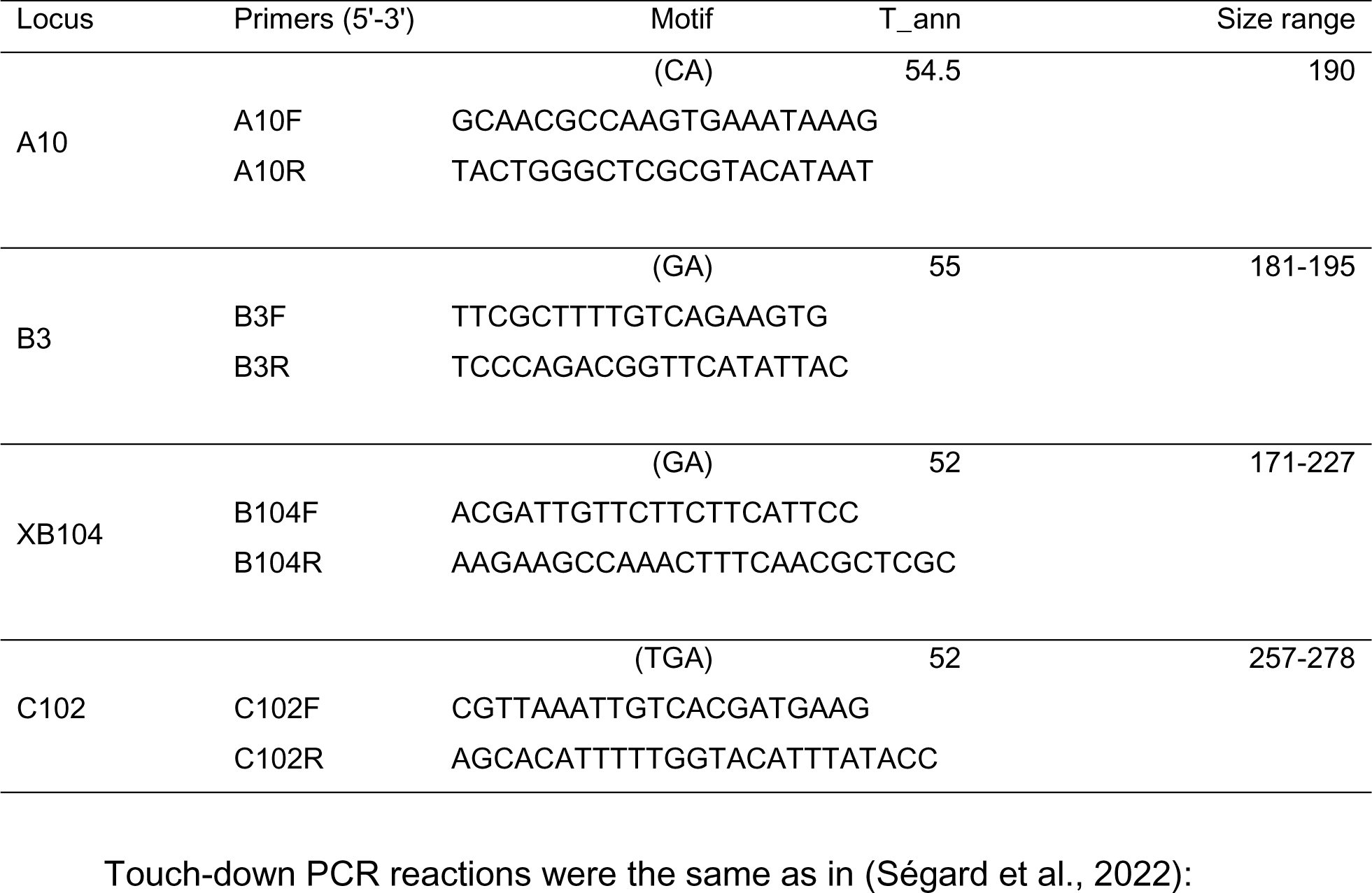
Information on microsatellite loci provided by A. Robinson (Insect Pest Control Sub-program, Joint Food and Agriculture Organization of the United Nations/International Atomic Energy Agency Program of Nuclear Techniques in Food and Agriculture) used in the present study. Name of primers motif, annealing temperature (T_ann in °C), the range of sizes of amplified fragments, and primers’ sequences are indicated.

